# Sepsis induces long-term reprogramming of human HSPCs and drives myeloid dysregulation in sepsis survivors

**DOI:** 10.1101/2024.12.14.628447

**Authors:** Marco De Zuani, Petra Lázničková, Marcela Hortová Kohoutková, Veronika Bosáková, Ivana Andrejčinová, Natália Vadovičová, Veronika Tomášková, Alexandra Mýtniková, Julie Štíchová, Tomáš Tomáš, Jiří Hrdý, Kristýna Boráková, Stjepan Uldrijan, Marcela Vlková, Vladimír Šrámek, Martin Helán, Kamila Bendíčková, Jan Frič

**Affiliations:** International Clinical Research Center, St. Anne’s University Hospital, Brno, Czech Republic; International Clinical Research Center, Faculty of Medicine, Masaryk University, Brno, Czech Republic; Department of Biology, Faculty of Medicine, Masaryk University, Brno, Czech Republic; Department of Anaesthesiology and Intensive Care, St. Anne’s University Hospital and Faculty of Medicine, Masaryk University, Brno, Czech Republic; Institute of Clinical Immunology and Allergology, St. Anne’s University Hospital and Faculty of Medicine, Masaryk University, Brno, Czech Republic; First Department of Orthopaedic Surgery, St. Anne’s University Hospital and Faculty of Medicine, Masaryk University, Brno, Czech Republic; Institute of Clinical Immunology and Allergology, First Faculty of Medicine, Charles University and General University Hospital in Prague, Prague, Czech Republic; Neonatology Department, Institute for the Care of Mother and Child, Prague, Czech Republic; Department of Modern Immunotherapy, Institute of Hematology and Blood Transfusion, Prague, Czech Republic

## Abstract

Sepsis is a life-threatening condition characterised by an overwhelming immune response and high fatality. While most research has focused on its acute phase, many sepsis survivors remain immunologically weakened leaving them susceptible to serious complications from even mild infections. The mechanisms underlying this prolonged immune dysregulation remain unclear, limiting effective interventions. Here, we analysed whether sepsis induced long-term “training” in hematopoietic stem and progenitor cells (HSPCs), imprinting changes that persist in their myeloid progeny. Peripheral blood analysis of 8 sepsis survivors, 12 patients with septic shock, and 10 healthy donors revealed a significant expansion of CD38+ progenitors in survivors, with increases in megakaryocyte-erythroid and granulocyte-monocyte progenitors, and reduced mature neutrophil counts. This shift suggests impaired granulopoiesis, favouring immature, immunosuppressive granulocytes. Differentiated macrophages from survivors’ HSPCs exhibited impaired metabolic pathways after lipopolysaccharide stimulation, with downregulation of tricarboxylic acid cycle and glycolysis genes, indicating altered immune metabolism. Pathway analysis revealed enhanced type-I interferon (IFN) and JAK-STAT signalling in survivors’ macrophages, reflective of potentially tolerance-prone reprogramming. Finally, exposing healthy donor HSPCs to IFNβ during macrophage differentiation reduced HSPC proliferation, increased apoptosis, and induced a metabolic shift towards glycolysis over mitochondrial respiration. Together, these findings suggest that sepsis induces lasting reprogramming in HSPCs leading to myeloid progeny with altered immune memory that might drive immune dysregulation in survivors. These data open avenues to explore potential targets to better manage long-term immune alterations in sepsis survivors.

**KEY POINTS:**

- Sepsis induces long-term alterations in HSPCs, leading to the expansion of immature progenitors and metabolic dysregulation of their progeny.
- Type-I IFN signalling reprograms macrophage differentiation, affecting their metabolic function and reducing cell proliferation.

## INTRODUCTION

Sepsis is a life-threatening condition characterised by a dysregulated host response to infection, leading to organ dysfunction and often high mortality.^1–3^ In 2017, 48.9 million people were affected by this syndrome, with 19.7% succumbing to it.^4^ Survivors face long-term consequences, including increased susceptibility to secondary infections, persistent immune-cell alterations, low-grade inflammation, and release of damage-associated molecular pattern molecules.^5^ Mechanistically, monocytes from sepsis survivors undergo inflammatory reprogramming, which drives chronic inflammation and persistent immune activation. This process is sustained by elevated cytokines and nucleotide oligomerization domain-like receptor protein 3 (NLRP3) components that can persist for years after sepsis.^6^

Hematopoietic stem cells (HSCs), which are produced in the bone marrow (BM), give rise to all blood-cell lineages.^7^ In normal aging, HSCs frequency in the BM increases, with a tendency towards a myeloid bias.^8^ Interestingly, sepsis can similarly promote myelopoiesis and trigger emergency haematopoiesis, which eventually leads to accelerated immunosenescence.^5^ Prolonged myelopoiesis, evidenced by an expansion of myeloid-derived suppressor cells (MDSCs) and subsequent long-term changes in myeloid functions, is observed in sepsis survivors months after the acute phase.^9^ Work conducted in murine model of sepsis showed that after acute sepsis, HSCs are less responsive to granulocyte-colony stimulating factor and thus fail to induce granulopoiesis, suggesting HSCs exhaustion after severe infection.^10^ However, the mechanisms by which these changes impact long-term immune function in sepsis survivors remain unclear.

Trained immunity – or innate immune memory – is a phenomenon whereby cells of the innate immune system, such as monocytes and macrophages, undergo epigenetic reprogramming after exposure to certain pathogens, enhancing their response to future infection.^11^ In a process referred to as ‘central trained immunity’, HSCs can also acquire “memory”, generating myeloid progeny that retain this trained state.^12,13^ For example, Bacillus Calmette-Guérin (BCG) vaccination instructs murine HSCs to produce trained myeloid progeny, which subsequently protects against *M. tuberculosis* infection.^14^ In the context of sepsis, hematopoietic stem and progenitor cells (HSPCs) and BM-derived macrophages in surviving mice undergo epigenetic reprogramming,^11^ resulting in impaired cytokine production.^15^ Interestingly, metabolic changes, such as a shift toward glycolysis, are highly linked to the induction of these epigenetic modifications.^11^ For example, secondary lipopolysaccharide (LPS) challenges in mice treated with β-glucan (a trained immunity inducer in myeloid cells) leads to increases in myelopoiesis, glycolysis, and cholesterol biosynthesis.^16^

Although the impaired function of myeloid cells in sepsis survivors has been already reported, the signalling pathways and metabolic mechanisms underlying these long-term alterations remain unexplored. As such, we lack the tools to identify or modulate immune dysfunction in affected patients. Moreover, the long-term consequences of HSPC training during the acute phase of sepsis on immune resilience in survivors are unclear. To address these knowledge gaps, we compared the profile of circulating HSPCs and peripheral blood subpopulations in patients with acute septic shock, long-term survivors (Surv; average 13.5 months post-sepsis), and age-matched healthy donors (HD). We then assessed the transcriptional and functional profiles of HSC-derived macrophages from Surv and HD to pinpoint potential contributors to macrophage functional and metabolic impairments.

## METHODS

### Study participants

Twelve adult patients admitted to the intensive care unit (ICU) at St. Anne’s University Hospital in Brno (Czech Republic) with early septic shock were prospectively enrolled into the study cohort. Two patients died before reaching the second collection time point, while the first time point collection for another patient failed. Patients with chronic immunosuppression, ongoing active oncological disease, or who had received antibiotic therapy for more than 2 days were excluded. Additionally, eight sepsis survivors (Surv, 8 to 26 months - 13.5 months on average after the initial ICU admission) were retrospectively enrolled in the “sepsis survivor cohort”. Finally, 10 healthy age- and comorbidity-matched individuals were recruited into the “age-matched healthy donors” (HD) cohort at the First Department of Orthopaedic Surgery, St. Anne’s University Hospital in Brno. Patients with an acute infection within the last 28 days, ongoing active oncological disease, or chronic immunosuppression were not included. Cohort details are summarised in Supplementary Tables 1 and 2. For *in vitro* studies on the effect of IFNβ on HSPCs differentiation, buffy coats from adult blood donors were obtained from the Department of Transfusion & Tissue Medicine of Brno University Hospital. Cord blood was obtained from women after childbirth at the Institute for the Care of Mother and Child in Prague. Written informed consent was obtained from all enrolled patients. All procedures were approved by the institutional Ethical Committee of St. Anne’s University Hospital Brno (4G/2018; 10G/2021), Ethical Committee of the Faculty of Medicine of Masaryk University (18/2023), and Ethical Committee of the Institute for the Care of Mother and Child (31/03/2014). All procedures complied with the Helsinki Declaration of 1975, as revised in 2013.

### Flow cytometry

Flow cytometry analyses followed the guidelines by Cossarizza et al.^17^ A total of 200 µL and 500 µL of heparinized blood were used to label mature immune cells and circulating HSPCs, respectively. Whole blood was lysed in 1x RBC lysis buffer for 10 minutes at room temperature (RT). Where indicated, dead cells were labelled with Live/Dead fixable dyes (Thermo Fisher Scientific) at a concentration of 1:800 in PBS. Cells were labelled in FACS buffer for 30 minutes on ice with the antibodies listed in Supplementary Table 3. Where indicated, propidium iodide was added immediately before sample acquisition to discriminate dead cells. Then, 10 µl of Precision Count Beads (BioLegend) were added to each sample to obtain absolute counts of HSPCs and other immune subsets. All samples were acquired on a Sony SA3800 spectral analyser (Sony Biotechnologies).

### HSPC-derived macrophage (HSDM) differentiation

HSDM differentiation was performed as described, with minor modifications.^18^ CD34+ cells were isolated directly from PBMCs or from enriched HSPCs (RosetteSep Hematopoietic Progenitor Enrichment Cocktail Kit, Stemcell Technologies) via immunomagnetic isolation (Miltenyi Biotec). Extended experimental details are reported in the supplementary methods.

### HSDM stimulation and RNA-sequencing

Mature HSDMs (1 x 10^5^) from HD and Surv were seeded in 96-well plates in Media C and stimulated with 100 ng/mL LPS-EB (Invivogen) for 3 hours at 37°C. Total RNA was extracted using the RNeasy Plus Micro Kit (Qiagen), according to manufacturer’s recommendations. RNA quality was assessed with Bioanalyzer2100 RNA Nano 6000 chips (Agilent Technologies), and samples with an RNA Integrity Number (RIN) > 8 were used for sequencing. An Illumina sequencing library was prepared using the NEBNext Ultra II Directional RNA Library Prep Kit (New England Biolabs) following the manufacturer’s protocols. Total RNA was used for poly-A enrichment, then fragmented, and reverse transcribed into cDNA. After universal adapter ligation, samples were barcoded using NEB dual indexing primers and pooled equimolarly after quantitation with PicoGreen. The sample pool was sequenced using a Nextseq 550 sequencer (Illumina) with a 75-cycle high-output cartridge.

### RNA-seq analysis

Raw reads were quality checked, pre-processed, and mapped to the reference genome (Ensembl GRCh38) with gene annotation (Ensembl v94). Mapped reads were counted and summarised by gene. After removing genes with < 10 counts, differentially expressed genes (DEGs) were calculated using DESeq2.^19^ Gene ontology (GO) and Gene Set Enrichment Analysis (GSEA) were performed using the clusterProfiler package.^20^ GSEA was performed after adaptive shrinkage of the log2 fold-change (LFC) values.^21^ To infer signalling pathway activity, the DecoupleR package^22^ was used, fitting a Multivariate Linear Model using as input the Wald statistic results from DESeq2 and the top 500 responsive genes ranked by p-value in the PROGENy collection^23^. Transcription factor activity was inferred by fitting a Univariate Linear Model (available with the DecoupleR package) using as input the Wald statistic results calculated by DESeq2 and the human regulons in the DoRothEA gene regulatory network with confidence levels “curated/high (A)”, “likely (B)”, and “medium (C)”.^24^

### Statistical analyses

Statistical analyses were performed with R v4.0.2. Specific statistical tests are reported in each figure legend. The Shapiro-Wilk test and visual inspection of QQ-plots were used to determine the normality of distributions, guiding the selection of parametric or non-parametric tests.

Sample collection and preparation, extended HSPC-derived macrophage (HSDM) differentiation, Immunofluorescence staining, Cell cycle profiling, Apoptosis assay, and Metabolic profiling of HSDMs are described in Supplementary methods and figures (Appendix 1).

## RESULTS

### Circulating HSPCs are expanded in sepsis survivors

We first aimed to investigate the effect of sepsis on HSPCs and their progeny, in order to understand whether the long-term immunosuppression observed in sepsis survivors could be due to the reprogramming of HSPCs during sepsis. To do so, we enrolled 12 patients with septic shock at two time points (within 24 hours (T1) or 3-5 days (T2) from ICU admission), eight Surv (average 13.5 months since ICU discharge; Supp. Table 1), and 10 HD (Supp. Table 2). We began by analysing HSPCs in the peripheral blood of all participants by flow cytometry (see Supp. Figure 1 for gating strategy) and found differences in their absolute numbers across groups (Figure 1A, Supp. Figure 2A).

**Figure 1.**
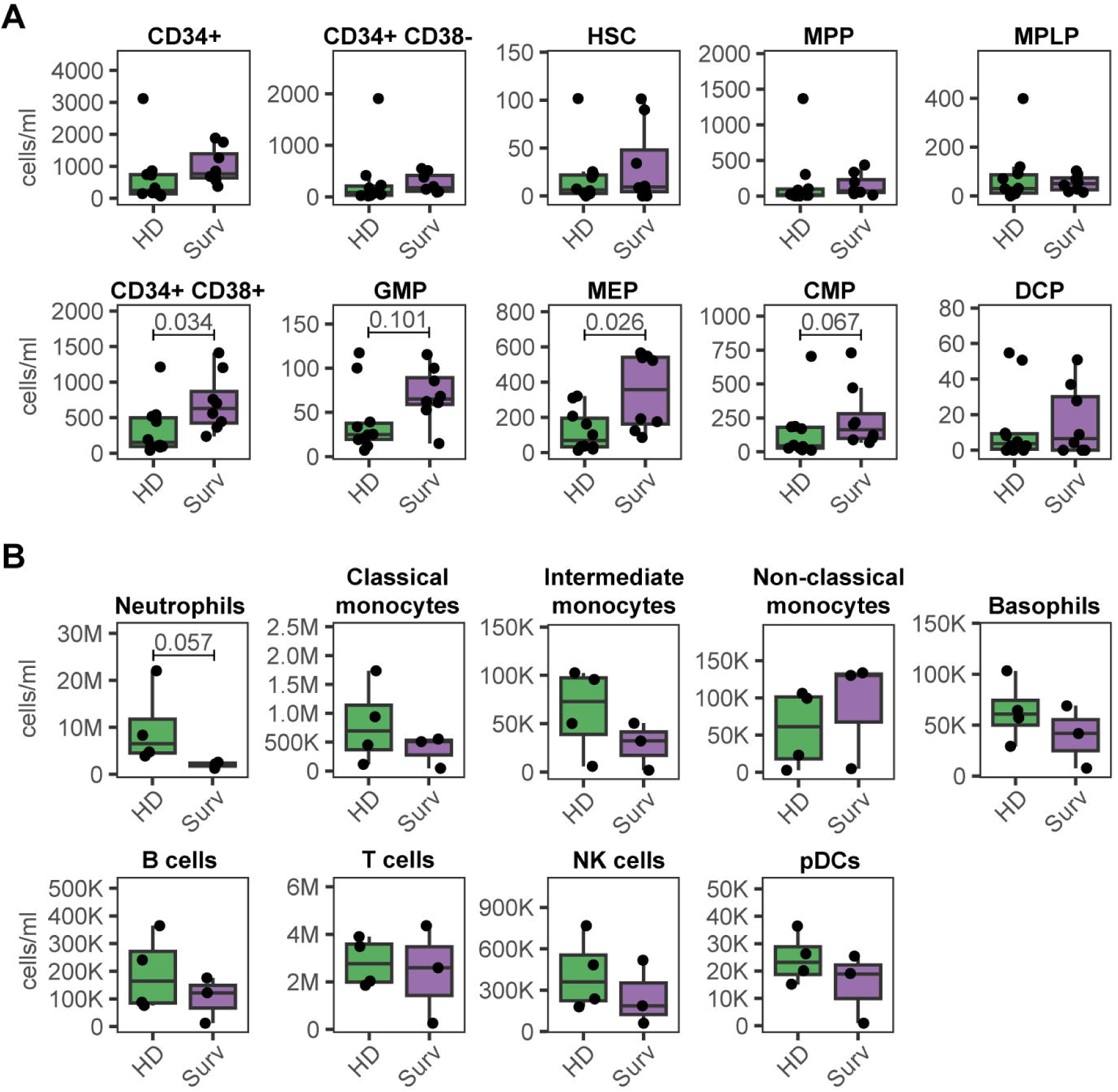
Flow cytometry phenotyping of HSPCs and mature immune cells in peripheral blood of Surv and HD. **A.** Box plots representing the absolute number of circulating HSPC subsets in age-matched healthy donors (HD) and long-term sepsis survivors (Surv). **B.** Box plots representing the absolute number of circulating immune cells in HD and Surv. The differences between the two groups were tested with a Wilcoxon rank-sum test. * = p-value ≤ 0.05.

When comparing HD with Surv, the latter showed a significant expansion of CD38+ progenitors (Figure 1A, p=0.034). Among these committed progenitors, megakaryocyte-erythroid progenitors (MEPs) were significantly increased in survivors, while common myeloid progenitors (CMPs) and granulocyte-monocyte progenitors (GMPs) showed a similar trend (Figure 1A, p=0.068 and p=0.101, respectively). When patients with septic shock were included in the comparison, we found a significant increase in the absolute GMP counts in survivors compared to septic shock patients at both time points (p=0.0276 for TP1 and p=0.0288 for TP2; Supp. Figure 2A). MEP counts showed a similar trend when compared to TP1 (p=0.0528). Taken together, these results suggest that septic shock affects the differentiation of HSCs into committed progenitors long after infection resolution, favouring the accumulation of MEPs and GMPs.

To determine whether the changes observed in the HSPC compartment were reflected in the terminal differentiation of blood cells, we analysed the peripheral blood of Surv. Compared to HD, Surv showed a decrease in absolute count of mature neutrophils (p=0.057, Figure 1B). Consistent with the expansion of polymorphonuclear (PMN) cells and early MDSCs in the peripheral blood of Surv,^9^ these findings suggest that sepsis induces long-term reprogramming of the hematopoietic compartment, promoting a skew towards the granulocytic lineage and the development of immature and immunosuppressive granulocytes. The release of MEP from BM in peripheral blood has been described in a mouse model of sepsis as a result of higher concentration of SCF in peripheral blood compared to BM.^26^ Here we report expansion of MEP in Surv long after septic shock, suggesting that a similar mechanism might be still taking place long after recovering from sepsis.

### Macrophages derived from sepsis survivor HSPCs show metabolic impairments

Having shown expanded CD38+ progenitors, increasing trends in CMP and GMP and decrease in absolute counts in neutrophils in Surv compared to HD, we hypothesized that HSPCs in Surv could give rise to a myeloid progeny with altered functionality. To test this, we adapted a protocol^18^ to differentiate macrophages *in vitro* from circulating CD34+ HSPCs (HSPC-derived macrophages, HSDMs). We first used flow cytometry to characterise mature HSDMs obtained from adult circulating HSPCs isolated from adult blood donors. These cells expressed CD68, CD11b, CD33, CD14, CD16, CD206, CD86, HLA-DR, CCR5, TLR2 and TLR4 (Supp. Figure 2B and C), indicative of successfully differentiated macrophages. Moreover, these cells demonstrated the ability to phagocytose and sequester *Staphylococcus aureus* within their lysosomes, as shown by the uptake of pHrodo *S. aureus* particles (Supp. Figure 2D), confirming that their core immune functionality is retained.

Next, we assessed whether the transcriptional profile of HSDMs differentiated from Surv and HD differed as a result of sepsis. Here, HSDMs from Surv and HD were stimulated with LPS and subjected to bulk RNA-seq. We found that the stimulated HSDMs from Surv exhibited upregulation of genes involved in the “response to lipopolysaccharide” and “regulation of innate immune response” pathways when compared to non-treated cells from the same individuals (Figure 2A and B).

**Figure 2.**
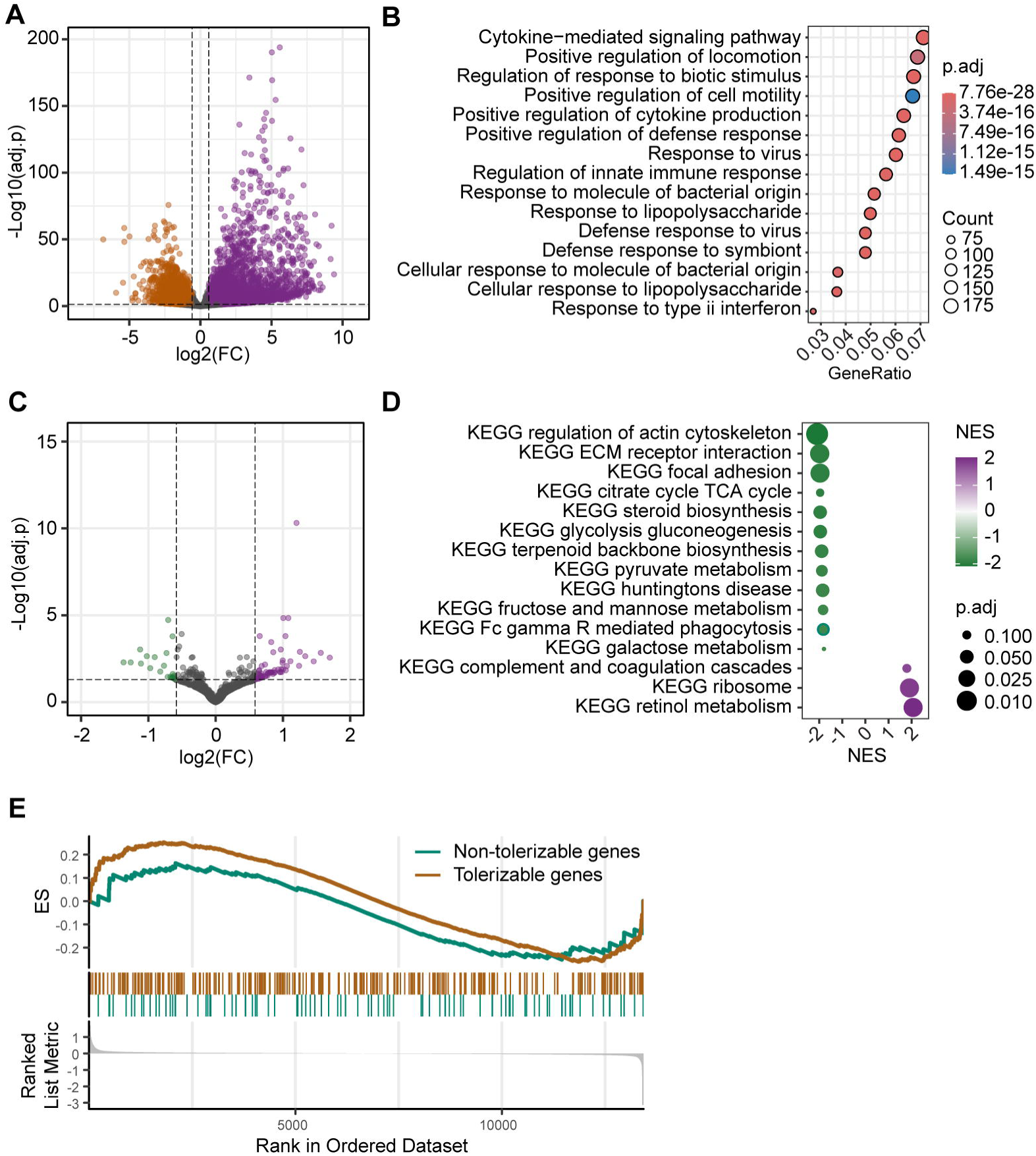
Transcriptional profiling of HSDMs from Surv and HD. **A.** Volcano plot comparing LPS-treated HSDMs and untreated HSDMs from sepsis survivors (Surv). Coloured dots indicate significant upregulation in LPS-treated (purple, lfc > 1.5 and adjusted p-value ≤ 0.05) or untreated (orange, lfc < 1.5 and adjusted p-value ≤ 0.05) cells. **B.** Top 15 upregulated “biological process” pathways in Surv HSDMs stimulated with LPS compared to untreated cells from the same patients. The colour of each dot represents the Benjamini-Hochberg adjusted p-value, while the size of the dot represents the number of genes enriched in each pathway. **C.** Volcano plot comparing HSDMs derived from Surv compared to HD after LPS stimulation. Coloured dots indicate significant upregulation in Surv (purple, lfc > 1.5 and adjusted p-value ≤ 0.05) or HD (green, lfc < 1.5 and adjusted p-value ≤ 0.05). **D.** Top 15 enriched terms from GSEA on the KEGG database, using differentially expressed genes between LPS-stimulated HSDMs derived from Surv and HD. Dot size indicates the adjusted p-value, and dot colour indicates the normalised enrichment score (NES). Green dots indicate terms significantly enriched in HSDMs from HD (i. e. NES < 0), while purple dots indicate terms significantly enriched in HSDMs derived from Surv. **E.** GSEA plot illustrating the enrichment of “tolerizable” (orange line) and “non-tolerizable” (green line) genes28 in LPS-stimulated HSDMs derived from Surv compared to HD. The bottom portion of the plot shows the ranked positions of genes by differential expression, with upregulated genes concentrated on the left (positive rank) and downregulated genes on the right (negative rank).

When comparing LPS-stimulated HSDMs from Surv and HD, we found that the former downregulated many genes involved in key metabolic pathways, including the TCA cycle, glycolysis/gluconeogenesis, and pyruvate metabolism, as indicated by the negative gene-set enrichment scores for these pathways (Figure 2C and D). These findings align with other studies indicating that trained immunity relies on the metabolic reprogramming of myeloid cells, and with studies showing a decrease in all major metabolic pathways as a hallmark of tolerant monocytes.^11,27^

As initial exposure to bacterial molecules (LPS in particular) can render myeloid cells more tolerant to subsequent restimulation,^27,28^ we explored whether a similar mechanism might affect HSDMs derived from Surv. Using a publicly available dataset of genes that are “tolerizable” and “non-tolerizable” to TLR4-dependent LPS exposure in human macrophages,^28^ we performed gene set enrichment analysis (GSEA) and found no enrichment in either category (adjusted p value=0.484 and 0.604, respectively; Figure 2E). This finding suggests that the mechanisms inducing a tolerant phenotype in macrophages after LPS exposure differs from the one acting on HSPCs during sepsis. It is, thus, possible that the mechanisms governing endotoxin tolerance and sepsis-mediated immunosuppression rely on different processes, the latter likely involving a metabolic and epigenetic rewiring of HSPCs.

### Type-I IFN signalling characterises HSDMs in sepsis survivors

To explore signalling pathways differences that might underlie our observations regarding the metabolic and transcriptional reprogramming of HSDMs from Surv, we inferred the activity of signalling pathways and transcription factor (TF) regulons across conditions using our RNA-seq data. In HSDMs from Surv, LPS stimulation led to induction of the NF-κB pathway (Figure 3A). This was accompanied by the activation of TFs involved in the TLR4-mediated response to LPS, including NFKB1, RELA and RELB (Figure 3B), which are central to NF-κB signalling.^29,30^ Additionally, we observed JAK-STAT pathway activation in response to LPS stimulation. While NF-κB is rapidly activated through TLR4 upon LPS exposure, the JAK-STAT pathway can be activated in an autocrine manner following the induction of IFNβ expression, which is also triggered by TLR4 activation via LPS.^29,30^

**Figure 3.**
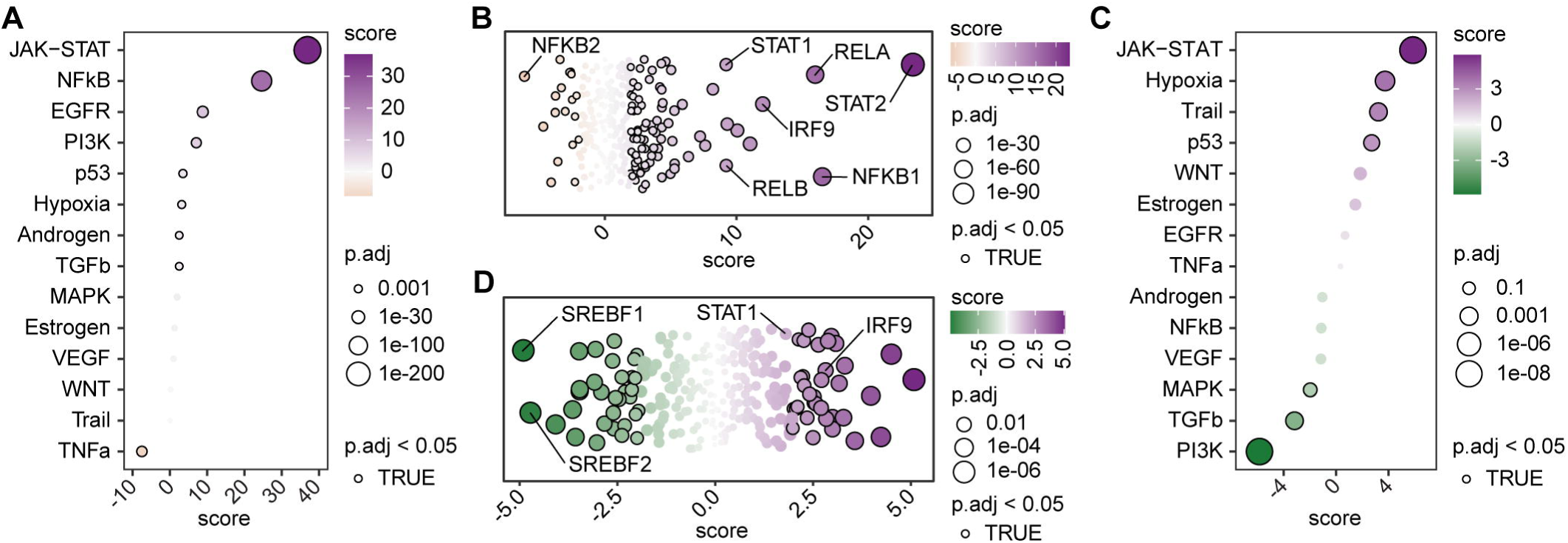
Signalling pathways and transcription factor activity in HSDMs from Surv and HD. **A.** Enriched signalling pathways in Surv HSDMs stimulated with LPS compared to untreated cells from the same donors. Dot size indicates the adjusted p-value, while dot colour indicates the enrichment score (purple, ES > 0 - enriched in LPS-treated HSDMs; orange, ES < 0 - enriched in untreated HSDMs). A black border around each dot indicates statistically significant enrichments (adjusted p-value ≤ 0.05). **B.** Transcription factor activity in Surv HSDMs stimulated with LPS compared to untreated cells from the same donors. Each dot represents a transcription factor, with dot size indicating the adjusted p-value, and dot colour indicating the activity score. Positive scores (purple) indicate increased transcription factor activity in LPS-treated HSDMs, while negative scores (orange) indicate increased transcription factor activity in untreated cells. A black border around each dot indicates statistically significant enrichments (adjusted p-value ≤ 0.05). **C.** Enriched signalling pathways in LPS-stimulated HSDMs derived from Surv versus HD. Dot size indicates the adjusted p-value, while dot colour indicates the enrichment score (purple, ES > 0 - enriched in Surv HSDMs; green, ES < 0 - enriched in HD HSDMs). A black border around each dot indicates statistically significant enrichments (adjusted p-value ≤ 0.05). **D.** Transcription factor activity in LPS-stimulated HSDMs derived from Surv versus HD. Each dot represents a transcription factor, with dot size indicating the adjusted p-value and dot colour indicating the activity score. Positive scores (purple) indicate increased transcription factor activity in Surv HSDMs, while negative scores (orange) indicate increased transcription factor activity in healthy donors’ HSDMs. A black border around each dot indicates statistically significant enrichments (adjusted p-value ≤ 0.05).

When directly comparing LPS-stimulated HSDMs derived from Surv with those from HD, we found distinct signalling patterns. Specifically, JAK-STAT pathway activity was higher in HSDMs from Surv, whereas the phosphatidylinositol 3-kinase (PI3K) pathway was more activated in HSDMs from HD (Figure 3C). Accordingly, we noticed the preferential activation of STAT1 and IRF9 in HSDMs from Surv, compared to those from HD (Figure 3D). These TFs are activated downstream of IFNAR1 and IFNAR2 receptors in response to type-I interferons (IFN) such as IFNα or IFNβ, and regulate the expression of IFN-stimulated genes (ISGs). Taken together, our results suggest a hyperactivation of type-I IFN signalling in macrophages differentiated from Surv HSPCs, which could reflect a mechanism to overcome long-term sepsis-related immunosuppression and tolerance.^31^

### IFNβ affects the development of macrophages from HSPCs

Given that HSDMs from Surv showed increased activation of the JAK-STAT pathway and elevated STAT1 and IRF9 TFs activity, we hypothesised that type-I IFNs could be responsible for the dysregulation observed in the HSPC compartment of Surv. To determine the role of type-I IFNs during myeloid differentiation, we differentiated HSDMs from circulating HSPCs isolated from adult healthy blood donors in the presence or absence (not treated, NT) of IFNβ, which is a primary mediator of type-I IFN responses that influences immune cell differentiation and inflammatory signalling.

We observed a reduction in total cell numbers after 7 and 14 days of differentiating IFNβ-stimulated HSPCs, compared to untreated cells (Figure 4A and B). This reduction was similarly observed when using cord-blood HSPCs (Supp. Figure 3A). Nevertheless, IFNβ-treated cells from adult blood donors showed an increased proportion of CD14+ cells at 14 days of differentiation (Supp. Figure 3B), suggesting that IFNβ stimulation might support monocytic differentiation over other lineages *in vitro*.

**Figure 4.**
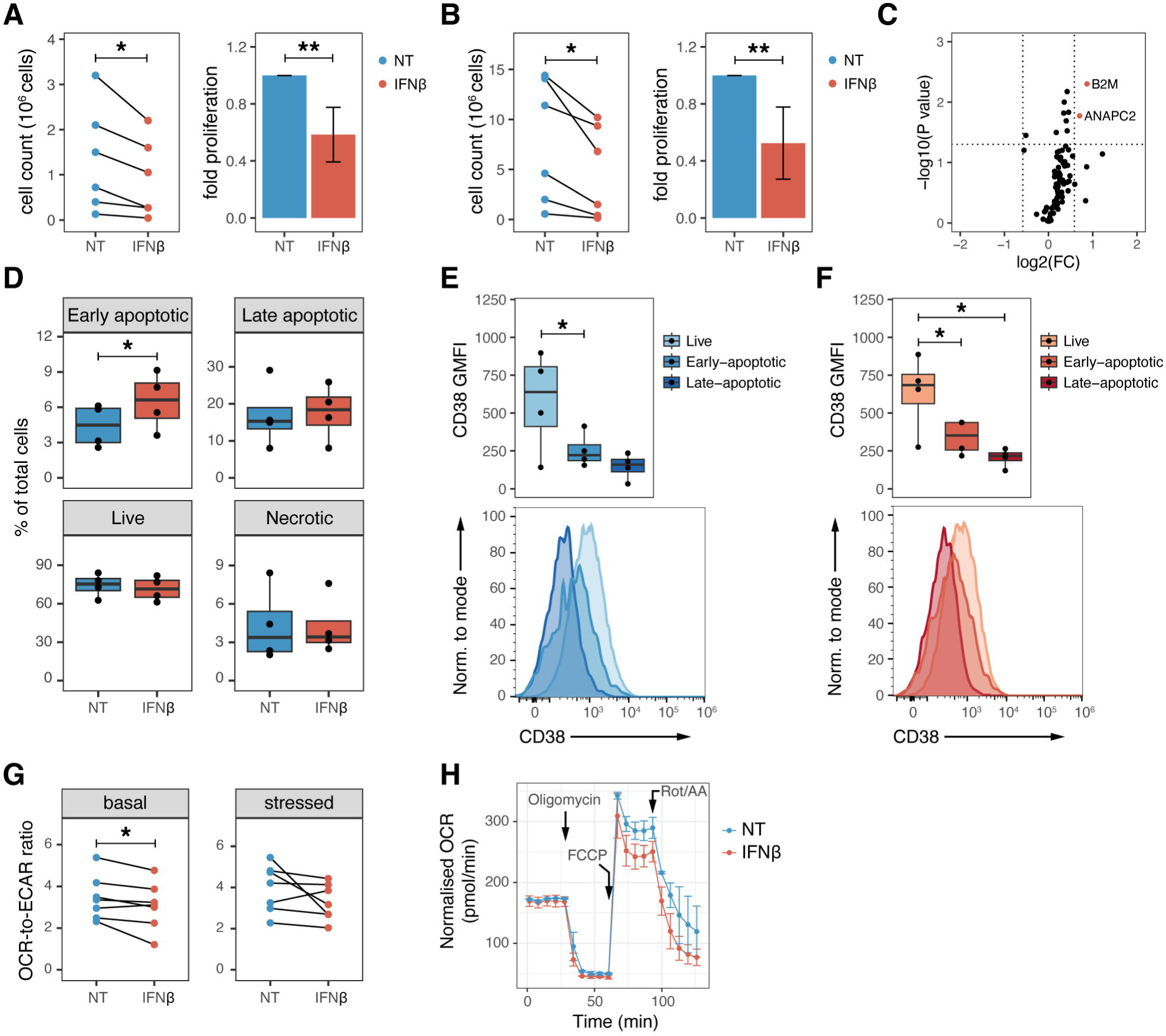
The effect of IFNβ on differentiation and function of HSDMs from adult blood donors. **A.** Comparison of cell counts after 7 days of differentiation of HSPCs from adult blood donors in the presence (red) or absence of IFNβ (blue). Left: total cell count, tested with a pairwise t-test. Right: fold-change compared to not treated (NT) cells from each donor, tested with a pairwise t-test. * = p-value ≤ 0.05, ** = p-value ≤ 0.01. **B.** Comparison of cell counts after 14 days of differentiation of HSPCs from adult blood donors in the presence (red) or absence of IFNβ (blue). Left: total cell count, tested with a pairwise t-test. Right: fold-change compared to non-treated cells from each donor, tested with a pairwise t-test. * = p-value ≤ 0.05, ** = p-value ≤ 0.01. **C.** Volcano plot comparing IFNβ-treated (red) and NT cells (blue). Dotted lines indicate statistical significance thresholds (|FC| ≥ 1.5, p-value ≤ 0.05). **D.** Percentage of early apoptotic, late apoptotic, live, and necrotic cells in NT (blue) and IFNβ-treated cells (red) after 7 days of differentiation. Differences between the two groups were tested with a pairwise t-test. * = p-value ≤ 0.05. **E.** Top: geometric mean fluorescence intensity (GMFI) of CD38 in live, early-apoptotic and late-apoptotic cells in NT samples. Differences between all groups was tested with a Tukey post-hoc test. * = p-value ≤ 0.05. Bottom: representative histogram showing CD38 levels in early-apoptotic and late-apoptotic cells in NT samples. **F.** Top: GMFI of CD38 in live, early-apoptotic and late-apoptotic cells in IFNβ-treated samples. Differences between all groups was tested with a Tukey post-hoc test. * = p-value ≤ 0.05. Bottom: representative histogram showing CD38 levels in early-apoptotic and late-apoptotic cells in IFNβ-treated samples. **G.** Dot plot comparing the OCR-to-ECAR ratio in resting (left) and stressed (right) cells differentiated in the presence (red) or absence (NT, blue) of IFNβ. Differences between the two groups were tested with a pairwise t-test. * = p-value ≤ 0.05. **H.** Graph depicting the oxygen consumption rate (OCR) in HSDM differentiated in the presence (red) or absence (NT, blue) of IFNβ at basal level, and after addition of oligomycin, FCCP, and a combination of rotenone and antimycin A (Rot/AA).

To assess whether the reduced cell numbers in IFNβ-stimulated HSPCs from adult healthy blood donors were due to decreased proliferation, we analysed a set of genes involved in cell proliferation after 7 days of differentiation. Only two genes (*B2M* and *ANAPC2*) were significantly upregulated in IFNβ-stimulated cells compared to untreated cells, indicating that cell proliferation rates likely remained unchanged (Figure 4C). We then tested whether increased cell death during differentiation could account for the lower cell counts. Annexin-V staining revealed a significantly higher frequency of early apoptotic cells (Annexin-V+, PI-) in IFNβ-treated cells (Figure 4D). Interestingly, early and late apoptotic cells showed reduced expression of CD38 compared to live cells (Figure 4E and F), suggesting that these cells might represent less committed progenitors. We speculate that the increase in CD34+CD38+ HSPCs and GMP in Surv compared to HD and the concomitant decrease in mature neutrophils in peripheral blood in Surv could be caused by the IFNβ-induced apoptosis of CD38+ progenitors (Figure 1, Supp. Figure 2A). This would also be supported by the higher activity in the p53 pathway in HSDMs from Surv compared to HD (Figure 3C).

Finally, we examined whether IFNβ stimulation during HSDM differentiation induced any metabolic reprogramming. IFNβ-stimulated HSDMs showed a significantly lower oxygen consumption rate-to-extracellular acidification rate (OCR-to-ECAR) ratio at basal levels (Figure 4G, H). This finding suggests that, in the resting state, IFNβ-stimulated HSDMs depend less on the TCA cycle and more on glycolysis, compared to untreated cells, which is consistent with a metabolic shift following classical macrophage activation.^32^ Nevertheless, in Surv the transcriptional regulation of the glycolytic pathway in LPS-stimulated HSDMs is also suppressed (Figure 2D), likely due to the immunosuppression induced by the cytokine storm and their consequential switch towards alternatively-activated macrophages, as evidenced elsewhere,^1^ despite the activated hypoxia pathway (Figure 3C) that in general favours glycolysis in pro-inflammatory macrophages.^33^

## DISCUSSION

Sepsis research has traditionally focused on immune-cell responses during the acute phase due to its high fatality rate in this period. However, the persistent health complications observed in many sepsis survivors months after their initial recovery, highlights a significant gap in our understanding of the long-term impacts of sepsis on the immune system. This study aimed to explore these long-term effects specifically on HSPCs, investigating whether sepsis induces “training” in these cells during septic shock that carries over to the myeloid progeny and alters their metabolism and signalling responses.

Previous studies, including our own,^9,34^ have identified alterations in immune cell phenotypes in sepsis patients during the acute phase of septic shock,^35–37^ as well as notable shifts in MDSCs and PMN-MDSCs that are detectable 6-26 months after recovery.^9^ Nevertheless, a thorough analysis of the immunophenotypic changes that are apparent in sepsis survivors has not yet been performed, thus limiting our understanding of the adverse effects and recurrence of sepsis in these patients. We now provide evidence of a likely biologically relevant decrease in absolute neutrophil numbers in the peripheral blood of long-term sepsis survivors (Surv). Furthermore, we observed an expansion of GMPs in Surv compared to patients in the acute phase of septic shock, indicative of defective granulopoiesis in this group. Together with an increase of immature PMN-MDSCs, these data suggest that impaired granulopoiesis gives rise to suppressor cells rather than mature neutrophils in sepsis survivors.

We also observed a long-term shift in the hematopoietic compartment of Surv, characterised by an increased number of committed CD38+ progenitors and MEPs compared to HD. Although thrombocytopenia commonly occurs in septic patients,^38^ thromboembolism events are also common.^39^ Accordingly, in murine models of severe septic shock it has been reported that MEP counts increase^26^ and that platelets can exacerbate inflammation^40^. The MEP expansion observed in Surv might explain the increased risks of myocardial infarction and stroke observed in these patients.^41^ The increase of CD38+ progenitors suggests that HSCs might undergo “training” during septic shock, which influences their differentiation and the characteristics of their myeloid progeny. Indeed, these findings resonate with an earlier finding that BCG vaccination trains monocyte and macrophage progeny.^14^ We confirmed whether there exists a memory of sepsis-related training in HSPC progeny by differentiating macrophages (HSDMs) *in vitro* from Surv HSPCs and re-challenging them with LPS. Bulk RNA sequencing revealed downregulation of genes involved in glycolysis/gluconeogenesis, the TCA cycle, and pyruvate metabolism in these LPS-challenged Surv HSDMs compared to HD. These findings align with previously described defects in glycolysis and oxidative phosphorylation in monocytes from septic patients.^27^ Despite published data suggesting normalisation of these metabolic determinants within days of recovery (> 7 days after sepsis),^27^ we show that such defects persist in HSDMs differentiated from Surv. These discrepancies might be caused by the different time frame after which the measurements were taken in these studies - quite early after the recovery vs months after the resolution of sepsis. We speculate, therefore, that sepsis primes HSPCs in a way that persists over time, potentially driving a myeloid bias in immune cell production.

LPS-induced tolerant macrophages can protect against septic-shock-induced cell death and improve survival in mice.^42^ Reprogramming into hyporesponsive, immunosuppressive phenotypes depend on the p21-mediated DNA binding of NF-κB p50 homodimers, which inhibit mRNA transcription and reduce IFNβ production in mice.^43^ Here, we demonstrate that reprogramming during septic shock differentially affects the response to LPS of HSDMs derived from Surv compared to HD HSDMs. Pathway analysis provided further insights into the altered immune responses in Surv as a result of this reprogramming: LPS stimulation induced NF-κB and JAK-STAT pathway activation in HSDMs from Surv, the JAK-STAT response notably stronger than observed in HD. This effect was coupled with elevated type-I IFN activity, including STAT1 and IRF9 activation that is indicative of sustained type-I IFN signalling. JAK-STAT activation after LPS treatment and IFNAR1 via autocrine IFNβ production are consistent with findings from murine BM-derived macrophages.^44^ The role of *Ifnar1* in immune “training” has also been described in murine alveolar macrophages,^45^ as well as in a mouse model of autoimmune systemic lupus erythematosus, where a type-I IFN signature promoted myelopoiesis.^46^ Additionally, type-I IFN signalling impairs macrophage anti-*Mycobacterium tuberculosis* immunity in mice.^47^ While IFNAR1 downregulation contributes to HSCs maintenance,^48^ heightened IFNβ signalling through JAK-STAT in Surv contrasts with the hyporesponsive phenotype seen in murine macrophages, where IFNβ production is suppressed.^43^ Together, these findings suggest that immune training in Surv is characterized by a long-lasting reprogramming of myeloid cells characterised by an immunosuppressive metabolic state and heightened type-I IFN responses upon LPS challenge. These results point towards a potential model where “primed” HSPCs differentiate into myeloid cells more prone to mounting antiviral over antibacterial responses, possibly underlying the susceptibility to bacterial infections of sepsis survivors and the potential sepsis recurrence.^49^

Compared to HSDMs derived from Surv, we found that HSDMs from HD predominantly activate the PI3K pathway upon LPS treatment. PI3K signalling is associated with modulating inflammation and enhanced cell survival, and so might constitute a protective mechanism to counterbalance excessive pro-inflammatory immune activation and cellular apoptosis.^50^ Interestingly, a dependency on PI3K has been shown in IFNβ-driven regulation of glucose metabolism.^51^ Our findings suggest that IFNβ exposure during HSPC differentiation from adult blood donors leads to increased early apoptosis (after 7 days of differentiation *in vitro*) and a bias towards CD14+ cell development (after 14 days of differentiation *in vitro*). Finally, we found that HSDMs differentiated in the presence of IFNβ depend on glycolysis. This finding is consistent with our RNAseq data, which revealed reduced TCA cycle activity in HSDMs derived from Surv but contradicts the reduced glycolysis observed in HSDMs from Surv. This is likely due to the different activation state of the cells and the length of the stimulation. A study evidenced that a 5-day “chronic” exposure of human monocyte-derived macrophages to IFNβ suppresses both basal oxygen consumption rate and glycolysis.^52^

Despite the limited number of samples analyzed and the simplified model of type-I IFN activity in sepsis modelled by prolonged IFNβ stimulation of HSPCs from adult blood donors, our study provides evidence of long-term reprogramming in Surv compared to HD. It is important to note that the cytokine storm and signalling dynamics during septic shock *in vivo* are greatly more complex than the *in vitro* system tested in our study. Moreover, different infectious agents and sepsis severity might be driving different responses, and thus reprogramming, in HSPCs. This could not be tested in our study due to the limited size of our cohort. Finally, our results highlight the need to evaluate the metabolic and transcriptional status of terminally differentiated immune cells during and after sepsis. These analyses could support the development of therapeutic targets to overcome the recurrence and adverse effects of septic shock.

Taken together, we show that sepsis induces a long-term skew in HSPC differentiation and myeloid cell functionality in Surv. Compared to HD, Surv show increased numbers of committed progenitors and decreased neutrophil counts. Importantly, the immune training of HSPCs during the acute phase of sepsis predisposes them to differential signalling upon subsequent LPS re-challenge *in vitro*. The evidence of metabolically impaired macrophages should be elaborated further and potentially used as a model to improve immune cell responses in sepsis survivors who suffer from opportunistic infections and face potential sepsis recurrence. Potential therapeutic strategies targeting immune cell-specific metabolic manipulations using nutrition supplements, or affecting the cytokine availability, such as IFNβ, could help re-establish the homeostatic state of hematopoietic progenitors and immune cells after the septic shock episode. Overall, our results open potential therapeutic opportunities to manage long-term immune-cell reprogramming in septic shock survivors.

## Supporting information

Appendix_1_Suppplementary Methods and Figures

## ACKNOWLEDGEMENTS

The research was supported by the Ministry of Health of the Czech Republic, grant nr. NV21J-05-00056), all rights reserved and DRO (Institute of Hematology and Blood Transfusion – UHKT, 00023736). The research was also supported by project nr. LX22NPO5107 (MEYS): Financed by European Union – Next Generation EU and the European Union’s Horizon Europe research and innovation programme under grant agreement No. 101137484.

We would like to thank the technical support team of the Center for Translational Medicine for technical support. Core Facility Genomics of CEITEC Masaryk University is gratefully acknowledged for the obtaining of the scientific data presented in this paper. We would also like to thank Dr. Jessica Tamanini from Insight Editing London for critical review of the manuscript.

## AUTHORSHIP

### Contribution

MDZ and JF designed the study; MH, VS, MV, and JH supervised the cohort recruitment; VT, AM, JS, TT, KrB, and MH recruited the study participants; MDZ, PL, MHK, KaB, VB, IA, NV, and SU performed the experiments and analyzed the data; MDZ, KaB, MHK, and JF secured funding; MDZ, PL, KaB, and JF wrote and reviewed the manuscript.

## APPENDIXES

Appendix 1: Supplementary Methods and Figures.

Appendix 2: Supplementary Tables.

## DATA AND CODE AVAILABILITY

The code used to analyse the bulk RNA-seq data and generate the figures is available at https://github.com/Deusu/HSDM-sepsis. The RNA-seq data generated in this study is available at Zenodo (10.5281/zenodo.14295543).

