## Appendix_1_Suppplementary Methods and Figures for "Sepsis induces long-term reprogramming of human HSPCs and drives myeloid dysregulation in sepsis survivors"

Running title: Sepsis induces long-term reprogramming of HSPCs

Marco De Zuani[1,#], Petra Lázničková[1,2,#], Marcela Hortová Kohoutková[1,2,*], Veronika Bosáková[1,2], Ivana Andrejčinová[1,2,3], Natália Vadovičová[1,3], Veronika Tomášková[4], Alexandra Mýtniková[1,4], Julie Štíchová[5], Tomáš Tomáš[6], Jiří Hrdý[7], Kristýna Boráková[8], Stjepan Uldrijan[1,3], Marcela Vlková[5], Vladimír Šrámek[4], Martin Helán[1,4], Kamila Bendíčková[1,2], Jan Frič[1,2,9,*]

*1 International Clinical Research Center, St. Anne's University Hospital, Brno, Czech Republic.*
*2 International Clinical Research Center, Faculty of Medicine, Masaryk University, Brno, Czech Republic.*
*3 Department of Biology, Faculty of Medicine, Masaryk University, Brno, Czech Republic.*
*4 Department of Anaesthesiology and Intensive Care, St. Anne's University Hospital and Faculty of Medicine, Masaryk University, Brno, Czech Republic.*
*5 Institute of Clinical Immunology and Allergology, St. Anne's University Hospital and Faculty of Medicine, Masaryk University, Brno, Czech Republic.*
*6 First Department of Orthopaedic Surgery, St. Anne's University Hospital and Faculty of Medicine, Masaryk University, Brno, Czech Republic.*
*7 Institute of Clinical Immunology and Allergology, First Faculty of Medicine, Charles University and General University Hospital in Prague, Prague, Czech Republic.*
*8 Neonatology Department, Institute for the Care of Mother and Child, Prague, Czech Republic*
*9 Department of Modern Immunotherapy, Institute of Hematology and Blood Transfusion, Prague, Czech Republic.*

*# These authors contributed equally to this work and share first authorship.*
*

Correspondence: Jan Frič, International Clinical Research Center, St. Anne's University Hospital Brno, Pekarska 53, Brno 60200, Czech Republic; e-mail address , Tel: +420 511 158 279.

### Supplementary methods

#### Sample collection and preparation

Blood samples were obtained from patients with septic shock, at two time points: within 12 hours (TP1), and at 3-5 days (TP2) after ICU admission. Blood samples from septic shock survivors were obtained at least 6 months (and up to 26 months) after hospital discharge. Buffy coats from adult blood donors were obtained from the Department of Transfusion &

Tissue Medicine of Brno University Hospital. Cord blood was obtained from women after childbirth at the Institute for the Care of Mother and Child in Prague. All samples were processed within 2 hours of collection. Blood was collected in BD Vacutainer® Tubes containing Sodium Heparin. Peripheral blood mononuclear cells (PBMCs) were isolated from 5 mL of heparinized blood and from buffy coats, by gradient centrifugation over Lymphoprep™ (1.077 g/mL, Alere Technologies AS) following manufacturer's recommendations. Cells were washed with fluorescence-activated cell sorting (FACS) buffer (PBS with 0.5% FBS and 2 mM EDTA) and immediately used for downstream analyses.

**HSPC-derived macrophage (HSDM) differentiation**

HSDM differentiation was performed as described, with minor modifications.[14] HSPCs were enriched from adult buffy coats or cord blood using the RosetteSep Hematopoietic Progenitor Enrichment Cocktail Kit (Stemcell Technologies), according to manufacturer’s recommendations. CD34+ cells were enriched with CD34+ Microbeads (Miltenyi Biotec), following the manufacturer’s recommendations. For sepsis survivors and HD, CD34+ cells were isolated directly from PBMCs. CD34+ cells were seeded at a concentration of 5 x 104 cells/cm2 in Media A (X-VIVO 15 supplemented with 50 ng/mL SCF, 15 ng/mL TPO, 30 ng/mL IL3, and 30 ng/mL Flt3-L). On day 3, media was changed by hemidepletion, without centrifuging the cells. On day 7, cells were collected, counted, and seeded at a concentration of 5 x 104 cells/cm2 in Media B (IMDM supplemented with 20% FBS, 1x GlutaMAX, 100 U/mL penicillin-streptomycin, 25 ng/mL SCF, 30 ng/mL M-CSF, 30 ng/mL IL3, and 30 ng/mL Flt3-L). On day 10, media was again changed by hemidepletion, without centrifuging the cells. On day 14, cells were enriched with CD14+ microbeads (Miltenyi Biotec) according to manufacturer’s recommendations. The CD14+ cells were seeded at a concentration of 2.5 x 105 cells/cm2 in Media C (RPMI supplemented with 10% FBS, 1x GlutaMAX, 100 U/mL penicillin-streptomycin, and 50 ng/mL M-CSF). On day 17, media was changed by hemidepletion, and on day 21, the cells were considered mature and used for downstream experiments. In some experiments, 50 ng/mL IFNβ (Stemcell Technologies) were added at the beginning of HSDM differentiation. HSDMs from Surv and HD were differentiated from CD34+ cells enriched directly from PBMCs using CD34+ Microbeads (Miltenyi Biotec), following the manufacturer’s recommendations.

**Immunofluorescence staining**

Mature HSDMs from HD were collected and resuspended at a concentration of $5 \times 10^5$ cells/mL in pre-warmed Media C. Then, 30 µl of cell suspension was applied to each channel of a µ-Slide IV 0.4 (Ibidi) and cells were allowed to attach for 2 hour at 37ºC. 5 µL of pHrodo™ Green *Staphylococus aureus* BioParticles™ Conjugate (Invitrogen) was added to each channel, incubated for 1 hour at 37ºC, and fixed with 4% paraformaldehyde in PBS for 20 minutes at RT. Cells were permeabilized with 0.2% Triton-X100 in PBS for 5 minutes at RT, stained with AlexaFluor-546-conjugated phalloidin (Invitrogen, 1:200 in PBS + 0.5% BSA) for 30 minutes at RT, and counterstained with DAPI. Images were acquired using a Zeiss LSM 780 confocal microscope.

**Cell cycle profiling**

Total RNA was extracted from 1 x $10^6$ HSDMs at day 7 of differentiation using the RNeasy Mini Plus Kit (QIAGEN) following the manufacturer's recommendations. Then, 500 ng of total RNA was reverse-transcribed using the RT2 Easy First Strand kit (QIAGEN). The resulting cDNA was used in the $RT^2$ Profiler™ PCR Array Human Cell Cycle (PAHS-020Z, QIAGEN), following manufacturer's recommendations. RT-qPCR was performed on a LightCycler 480 (Roche, Basel, Switzerland), and the results were analysed using QIAGEN's RT2 Profiler PCR Arrays & Assays Data Analysis software. Data was normalised against *GAPDH, RPLP0, HPRT1* and *ACTB.*

**Apoptosis assay**

CD34+ cells were isolated after 7 days of culture in Media A, in the presence or absence of 50 ng/mL of IFNβ and stained with the cocktail of antibodies indicated in Supplementary Table 3 ("apoptosis" panel), as described above. After staining, cell pellets were resuspended in 100 µL Annexin-V-Binding Buffer containing 5 µL of PE/Cy7-conjugated Annexin-V and 10 µL of propidium iodide solution (BioLegend), and incubated for 15 minutes at RT. Cells were analysed on a Sony SA3800 spectral analyser (Sony Biotechnology).

**Metabolic profiling of HSDMs**

After 21 days of differentiation, 0.8 x $10^5$ mature HSDMs were seeded in each well of a poly-L-lysine coated (Sigma-Aldrich) Seahorse XFp plate (Agilent Technologies) in HSDM Media C. One hour later, the media was replaced with Seahorse XF base medium supplemented with 1 mM Na-pyruvate (Sigma-Aldrich), 2 mM glutamine (Gibco) and 10 mM glucose (Sigma-Aldrich), and adjusted to pH 7.4. Cell metabolic profiles were assessed using the XF Cell Mito Stress Test Kit on a Seahorse XFp (Agilent technologies), following the manufacturer's instructions. Stressors were used at the following concentration: oligomycin 1 µM, FCCP 1 µM, Rotenone and Antimycin-A 0.5 µM. After each stressor injection, five cycles of measurement were taken. Cells in each well were fixed and permeabilized as described above, then stained with 1 µg/mL DAPI and imaged. CellProfiler[25] was used to enumerate the DAPI-positive nuclei in the central 10x field of each well, and the nuclei counts served to normalise the data. Data analysis was performed with the Wave software v 2.6.1.53 (Agilent

Technologies). Basal levels were determined at cycle 5, stressed levels at cycle 11 after FCCP injection.

### Supplementary Figures

#### Supplementary Figure 1

**A**

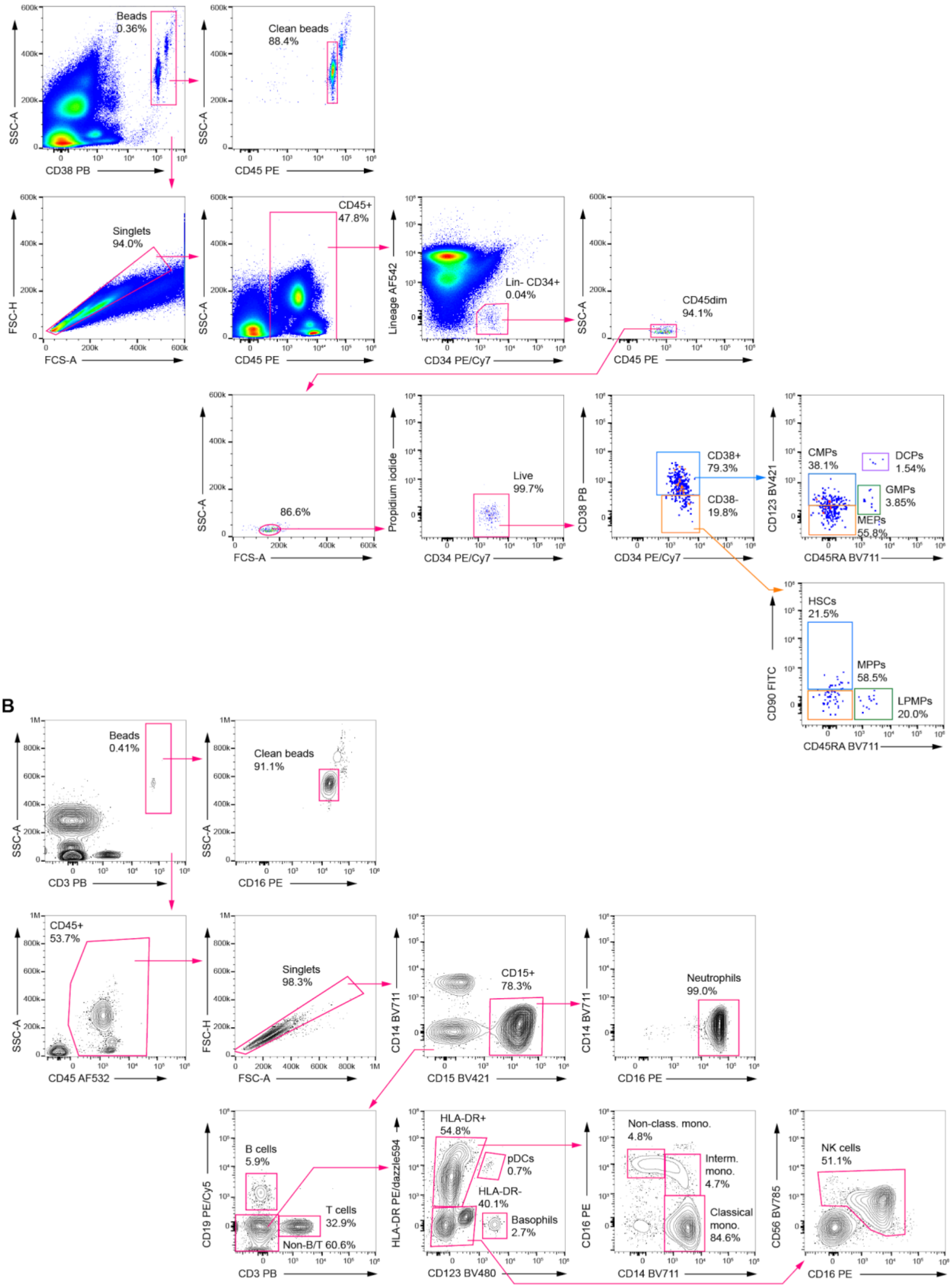

#### Supplementary Figure 2

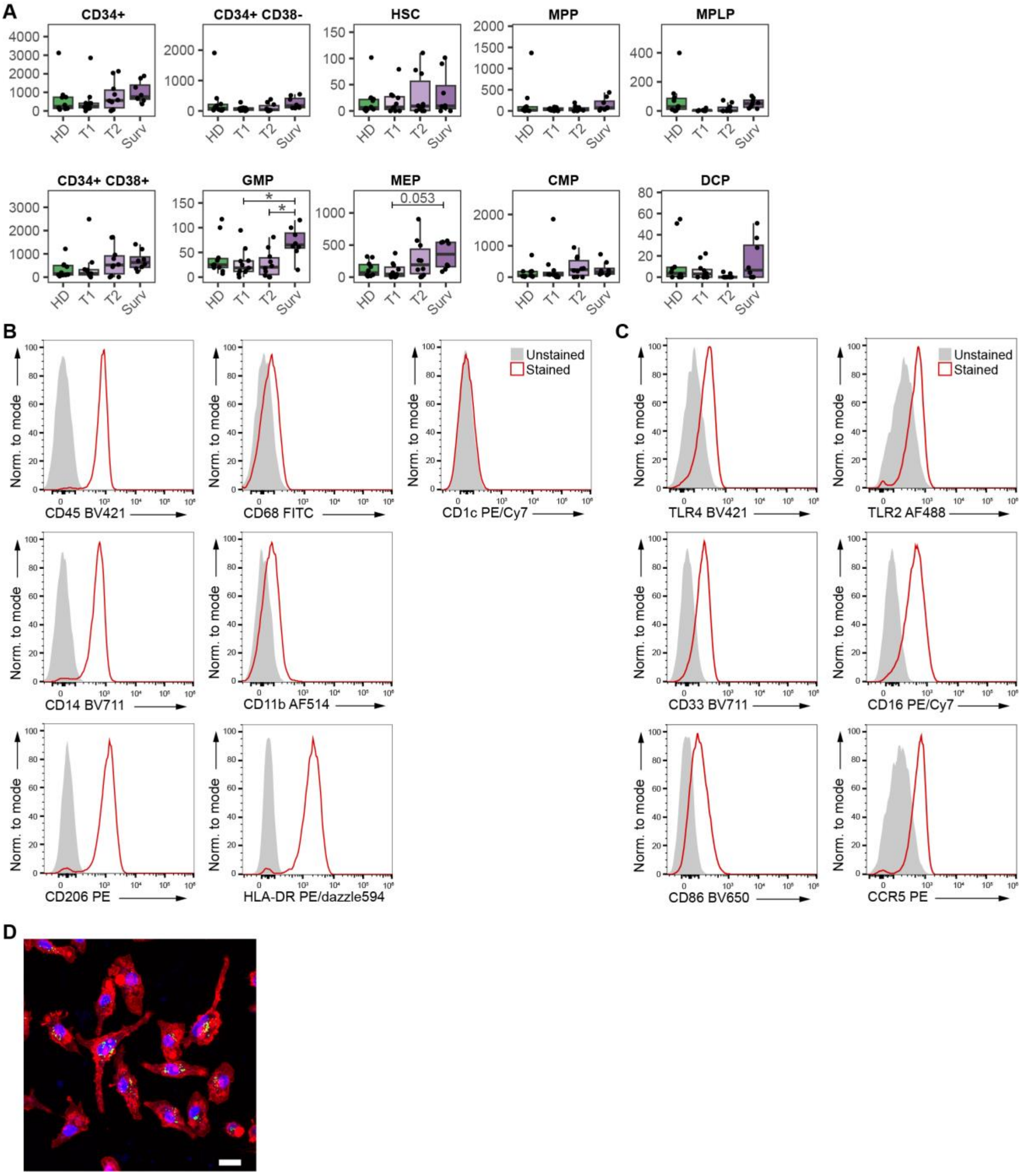

**Supplementary Figure 3**

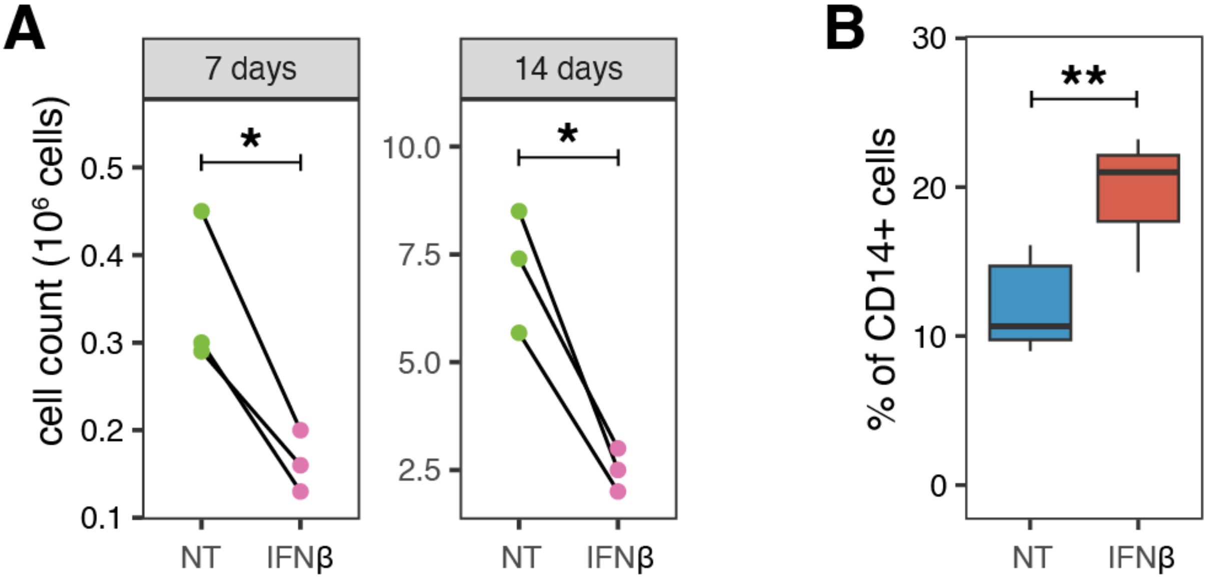

### Supplementary Figure Legend

**Supplementary Figure 1. Gating strategy for flow cytometry phenotyping of septic shock patients, Surv, and HD.**

**A.** Gating strategy to identify circulating hematopoietic stem cells and progenitors in blood of Surv and HD.

**B.** Gating strategy to identify the main leukocyte populations in whole blood of Surv and HD.

**Supplementary Figure 2. Flow cytometry phenotyping of HSPCs from septic shock patients, Surv, HD, and flow cytometry phenotyping and immunofluorescence staining of HSDMs from blood donors.**

**A.** Boxplots comparing the absolute number of circulating HSPC subsets in age-matched healthy donors (HD), septic shock patients at two time points (TP1, 12 hours post-ICU admission; TP2, 3-5 days post-ICU admission), and long-term sepsis survivors (Surv). Differences between all groups were tested using the Tukey post-hoc test. * = p-value ≤ 0.05.

**B.** and **C.** Histograms displaying the expression level of different myeloid cell markers on HSDMs from adult blood donors after 21 days of differentiation, as assessed by flow cytometry. Representative results of three independent experiments are shown.

**D.** Immunofluorescence staining of HSC-derived macrophages (HSDMs) from adult blood donors showing actin filaments (red, phalloidin), phagocytosed pHrodo™ Green Staphylococcus aureus BioParticles™ (green) and DAPI (blue). Scale bar: 10 µm.

**Supplementary Figure 3. Check-points of differentiation of HSDMs from cord blood and adult blood donors.**

**A.** Comparison of cell counts between untreated (light green) and IFNβ-treated (pink) cord-blood cells after 7 and 14 days of differentiation. Differences between the two groups were tested by pairwise t-test. * = p-value ≤ 0.05.

**B.** Percentage of CD14+ cells in untreated (blue) and IFNβ-treated (red) cells after 14 days of differentiation of HSPCs from adult blood donors. Differences between the two groups were tested by pairwise t-test. ** = p-value ≤ 0.01.

**Supplementary Table 1**

Acute sepsis patients and sepsis survivor's cohort information

**Supplementary Table 2**

Age-matched healthy donor cohort information

**Supplementary Table 3**

Antibodies, clones and dilutions used for the flow cytometry experiments performed in this study.
